# Strain-level overlap between infant and hospital fungal microbiomes revealed through *de novo* assembly of eukaryotic genomes from metagenomes

**DOI:** 10.1101/324566

**Authors:** Matthew R. Olm, Patrick T. West, Brandon Brooks, Brian A. Firek, Robyn Baker, Michael J. Morowitz, Jillian F. Banfield

**Affiliations:** department of Plant and Microbial Biology, University of California, Berkeley, CA, USA; Department of Surgery, University of Pittsburgh School of Medicine, Pittsburgh, PA, USA; Division of Newborn Medicine, Magee-Womens Hospital of UPMC, Pittsburgh, PA,USA; Department of Earth and Planetary Science, University of California, Berkeley, CA, USA; Department of Environmental Science, Policy, and Management, University of California, Berkeley, CA, USA; Earth Sciences Division, Lawrence Berkeley National Laboratory, Berkeley, CA, USA; Chan Zuckerberg Biohub, San Francisco, CA, USA

**Keywords:** Nosocomial infections, eukaryotes, metagenomics, genome-resolved metagenomics, strain-tracking, hospital microbiome, neonatal intensive care unit, premature infants

## Abstract

Eukaryotes are a leading cause of nosocomial infections in neonates, but their diversity and population heterogeneity are rarely investigated. This has led to an incomplete understanding of eukaryotic strains that colonize infants and of the neonatal intensive care unit (NICU) as a possible source of these strains. Analysis of 1,174 time-series metagenomes from 161 premature infants revealed fungal colonization of 13 infants, primarily in the first two weeks of life. Nearly all 24 NICU samples contained eukaryotes, and the most diverse communities were in NICU sinks. Five of fourteen newly-assembled eukaryotic genomes derived from genomically undescribed species. *Purpureocillium lilacinum* genomes from infant and NICU samples shared 99.999% average nucleotide identity, highlighting the potential of hospital-associated fungi to colonize hospitalized infants. We quantified zygosity and within-population variation associated with the diploid eukaryotes, and thus defined the genetic reservoirs of eukaryotes in room environments and infants.

## INTRODUCTION

Eukaryotes are common members of the human microbiome (Baley et al., 1986; Schulze and Sonnenborn, 2009). The colonization density and diversity of these eukaryotes are lower than their bacterial counterparts (Ott et al., 2008; Parfrey et al., 2011; Schulze and Sonnenborn, 2009), but they can have substantial health consequences. Certain eukaryotes can significantly reduce rates of antibiotic-associated diarrhea (Surawicz et al., 1989), limit bacterial populations through predation (Wildschutte et al., 2004), and are associated with asthma development in neonates (Fujimura et al., 2016). Neonatal nosocomial fungal infections have become more common with the increased use of invasive equipment in the neonatal intensive care unit (NICU) (Huang et al., 2000), and fungal organisms are now the third most common cause of sepsis in premature infants (Manzoni et al., 2007).

Fungal disease is most prevalent in premature infants and other immunocompromised patients (Fridkin and Jarvis, 1996), and several fungi causative of disease and their hospital reservoirs have been identified. For example, *Purpureocillium lilacinum* outbreaks have been traced to infected skin lotion (Orth et al., 1996) and neutralizing agents for artificial lenses (Pettit et al., 1980). In one case, fungal infections were traced to the water supply system of a paediatric institute (Mesquita-Rocha et al., 2013) and a NICU outbreak of the fungi *Malassezia pachydermatis* was linked to the dog of a healthcare worker (Chang et al., 1998). However, overall our understanding of the presence and diversity of fungi in hospital environments is lacking. Up to 20% of infections of medical implants are caused by *Candida albicans* (Nett and Andes, 2006), and several studies have linked *Candida parapsilosis* outbreaks to the hands of healthcare workers (Diekema et al., 1997; Huang et al., 1998a; Levin et al., 1998; Rangel-Frausto et al., 1999).

Most high-throughput studies of the hospital microbiome and the human gut microbiome use 16S rRNA gene sequencing, and thus are blind to eukaryotes because the primers do not target their 18S rRNA genes. Of five recent studies of the hospital microbiome, only one included primers to target the internal transcribed spacer (ITS) sequences to detect eukaryotes as well as bacteria (Bokulich et al., 2013; Hewitt et al., 2013; Lax et al., 2017; Oberauner et al., 2013; Shin et al., 2015). A recent review article referred to eukaryotes as a *“Missing Link in Gut Microbiome Studies”* (Laforest-Lapointe and Arrieta, 2018) and highlighted knowledge gaps related to the ecological roles, growth dynamics, and source of eukaryotes in the human and hospital microbiomes (Huffnagle and Noverr, 2013; Laforest-Lapointe and Arrieta, 2018).

An alternative approach to microbiome characterization involves shotgun metagenomics. In this method, all DNA from a sample is sequenced regardless of its organismal source or genetic context. New methods aid in reconstructing eukaryotic genomes from these datasets (West et al., 2018), enabling understanding of these organisms in the context of their entire communities, which also include bacteria, archaea, bacteriophage, viruses and plasmids. Relative to amplicon sequencing, genome assembly has several distinct advantages for understanding communities that contain eukaryotes. First, genomes provide information about *in situ* ploidy (number of distinct chromosomes sets per cell), heterozygosity (for diploid organisms, here used to refer to the fraction of alleles in a genome that have two versus one abundant sequence type), and extent of population heterogeneity (here used to refer to additional sequence types that constitute low abundance alleles). Second, strain-tracking can be performed using high-resolution genomic comparisons. Last, newly assembled eukaryotic sequences expand the diversity of genomically defined eukaryotes in public databases, enabling comparative and evolutionary studies.

Here, we used genome-resolved metagenomics to study eukaryote-containing microbiomes of premature infants and their NICU environment. We evaluated the incidence of eukaryotes in room and infant samples and investigated the time period during which infant microbiomes contained eukaryotes. Genomes were assembled for fourteen eukaryotic populations and their ploidy, zygosity, and population heterogeneity defined. Crucially, genomic identity was established for a fungus in an infant microbiome and the NICU environment, and a subset of other microbial eukaryotes in NICU rooms were classified as types that can cause nosocomial infections.

## RESULTS

### Recovery of novel eukaryotic genomes from metagenomes

In this study we analyzed 1,174 fecal metagenomes and 24 neonatal NICU metagenomes, totaling 5.31 terra-base pairs of DNA sequence (**Supplemental Table S1**). Fecal samples were collected from 161 premature infants primarily during the first 30 days of life (DOL) (full range of DOL 5 - 121; median 18), with an average of 7 samples per infant. NICU samples were taken from six patient rooms within the hospital housing the infants (Magee-Womens Hospital of UPMC, Pittsburgh, PA, USA). Three NICU locations were sampled in each room: swabs from frequently touched surfaces, wipes from other surfaces, and swabs from sinks (Brooks et al., 2017). Eukaryotic genomes were assembled from all samples using a EukRep based pipeline ((West et al., 2017); see methods section for details).

Fourteen novel eukaryotic genomes were recovered in total, with a median estimated completeness of 91% (Table 1). Genomes were assembled from organisms of a wide phylogenetic breadth, and five are the first genome sequence for their species (Figure 1). Twelve of the genomes are classified as fungal, and are described in more detail below. The two other genomes (both recovered from hospital sink samples) represent the first genomes of their phylogenetic Families. *Diptera S2_005_002R2* is within the phylogenetic clade of Diptera (true flies), and is equally related to *Drosophila melanogaster* (fruit fly) and *Lucila cuprina* (Australian sheep blowfly). *Rhabditida S2_005_001R2* is within the Family Rhabditida (nematode), and is related to both pathogenic and non-pathogenic roundworms. In both cases BLAST searches of the rpS3 protein sequence against NCBI revealed no significant hits, and furthermore, comparing the mitochondrial Cytochrome c oxidase subunit I (*COI*) gene and protein against the Barcode Of Life Database (BOLD) (Ratnasingham and Hebert, 2007) and NCBI also revealed no hits with high identity. Thus, we are unable to tie our genomes to any morphologically described species.

**Figure 1.**
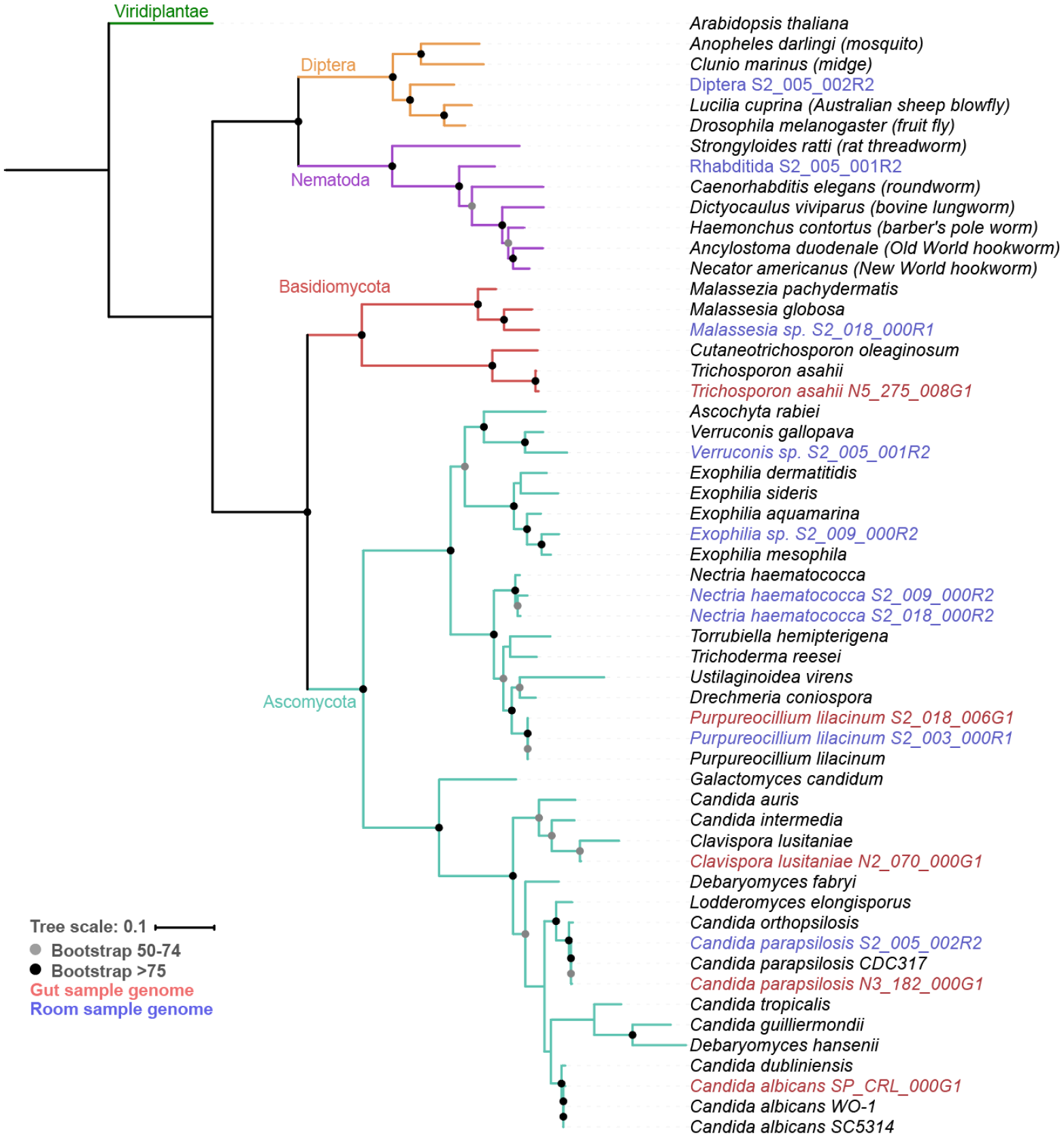
Phylogenetic tree of recovered eukaryote genomes. Genomes from infant-derived fecal samples (red), and NICU samples (blue) were phylogenetically placed using a phylogenetic tree based on the concatenation of the sequences of 16 ribosomal proteins (see methods). Branches with greater than 50% bootstrap support are labeled with their bootstrap support range. Reference ribosomal proteins were obtained from NCBI (NCBI Resource Coordinators, 2017) and the Candida Genome Database (Skrzypek et al., 2017).

**Table 1.** Description of *de novo* assembled eukaryotic genomes.

| Source | Genome | Comp. (%) | Length (bp) | N50 (bp) | Coverage |
| --- | --- | --- | --- | --- | --- |
| Infant gut | *Purpureocillium lilacinum* S2_018_006G1 | 98.4 | 35,688,710 | 422,361 | 20x |
| Infant gut | *Clavispora lusitaniae* N2_070_000G1 | 95.8 | 11,907,650 | 89,311 | 18x |
| Infant gut | *Candida parapsilosis* N3_182_000G1 | 96.7 | 12,563,647 | 65,710 | 182x |
| Infant gut | *Trichosporon asahii* N5_275_008G1 | 90.1 | 23,419,590 | 32,912 | 13x |
| Infant gut | *Candida albicans* SP_CRL_000G1 | 91.1 | 12,561,678 | 22,840 | 30x |
| NICU room | *Purpureocillium lilacinum* S2_003_000R1 | 98.4 | 35,724,498 | 520,486 | 67x |
| NICU room | *Malassezia* sp. S2_018_000R1 | 72.6 | 6,457,898 | 4,912 | 18x |
| NICU sink | *Nectria haematococca* S2_018_000R2 | 96.7 | 44,952,822 | 24,418 | 10x |
| NICU sink | *Candida parapsilosis* S2_005_002R2 | 92.8 | 11,573,959 | 14,507 | 9x |
| NICU sink | *Rhabditida* S2_005_001R2 | 74.9 | 50,505,025 | 8,214 | 8x |
| NICU sink | *Nectria haematococca* S2_009_000R2 | 73.6 | 31,143,909 | 8,000 | 7x |
| NICU sink | *Exophiala* sp. S2_009_000R2 | 75.9 | 24,670,482 | 7,386 | 7x |
| NICU sink | *Diptera* S2_005_002R2 | 52.5 | 43,769,201 | 6,834 | 10x |
| NICU sink | *Verruconis* sp. S2_005_001R2 | 52.8 | 15,639,153 | 5,112 | 6x |

### Fungal microbiome of the premature infant gut

Fungi were detected in 13 of the 161 premature infants profiled in this study (8%) (Figure 2A; **Supplemental Table S2**). Colonization tended to be sporadic, with relative abundance values shifting rapidly during infant development. Eukaryotes were detected significantly more often early in life, and significantly more often when antibiotics were recently administered (Figure 2). This is consistent with previous studies of fungal colonization of low birth weight infants (Baley et al., 1986; Huang et al., 2000).Interestingly, eukaryotic organisms in this study were also significantly anticorrelated with the abundance of bacteria of the phylum Proteobacteria (Figure 2B).

**Figure 2.**
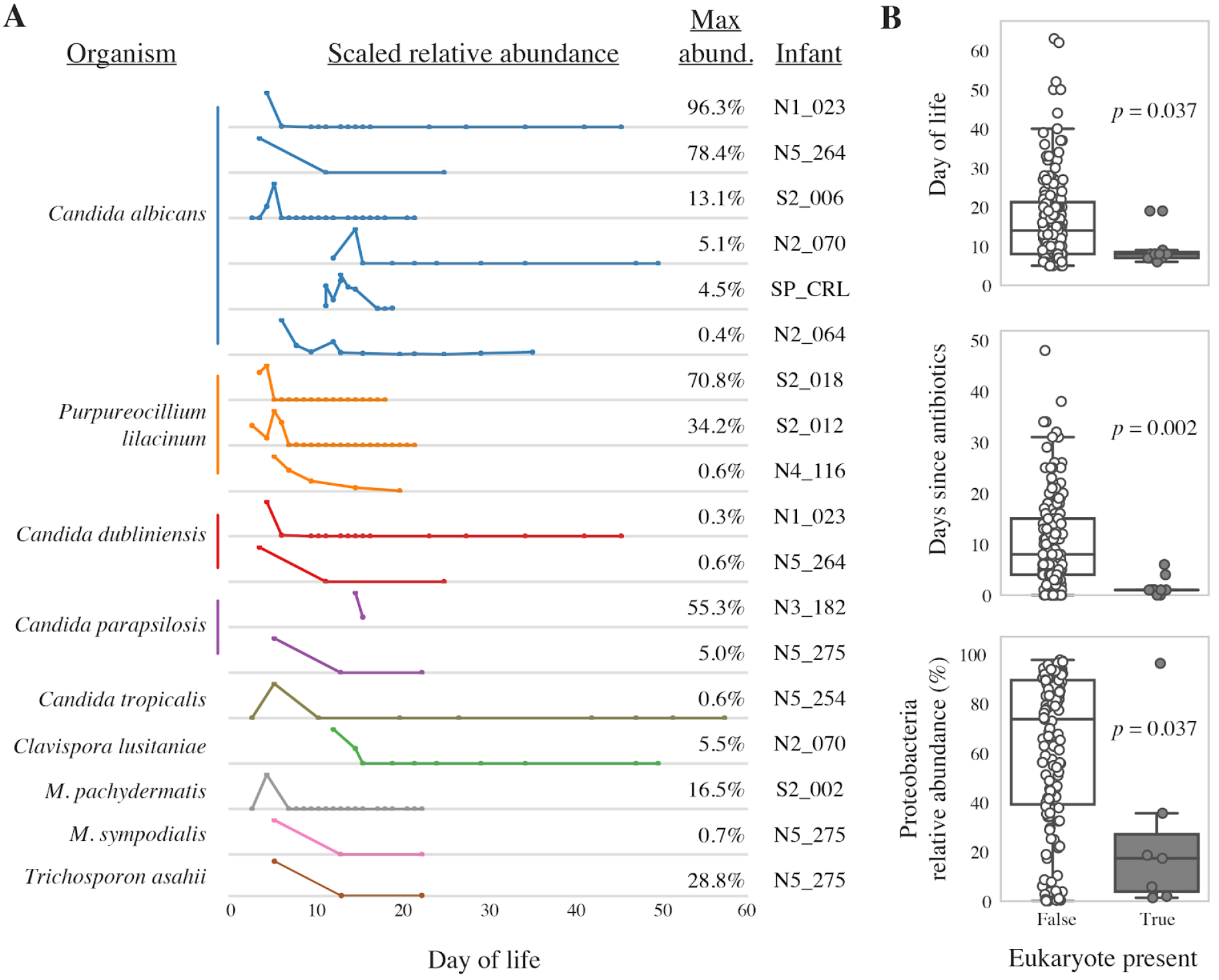
Abundance of eukaryotes colonizing infants. (A) The scaled relative abundance of each eukaryote in infants. Numbers on the right indicate the maximum relative abundance of the organism in that infant. The dots on the line-plots indicate the days of life on which fecal samples were collected and sequenced. (B) Metadata significantly associated with eukaryote abundance in infants. The distribution of values for all samples in which eukaryotes are not present (left; white box plot) compared to values of samples in which eukaryotes are present (right; grey box plot). The *p*-values were calculated using the Wilcoxon rank-sum test with Benjamini-Hochberg multiple testing *p*-value correction.

No infants colonized by eukaryotes in this study received antifungal antibiotics or showed any symptoms consistent with acute fungal infection. However, asymptomatic colonization has been shown to be a risk factor for fungemia (Huang et al., 1998b). Nine different eukaryotic species were detected in at least one infant, with *Candida albicans* and *Purpureocillium lilacinum* being the most common (Figure 2A). All nine species detected colonizing our premature infants have been previously implicated as agents of nosocomial infection (Table 2).

### Fungal microbiome of the neonatal intensive care unit

Eukaryotic organisms were detected in 22 of the 24 metagenomes of the NICU room environment (Figure 3). Eukaryotic DNA made up a median of 2.2%, 0.99%, and 0.08% of the communities in swabs, sink, and wipes, respectively. In order to compare the influence of room occupants and sampling location on the room mycobiome, we performed a non-metric dimensional scaling (NMDS) analysis (Figure 3A). Communities were strongly differentiated based on sampling location.

**Figure 3.**
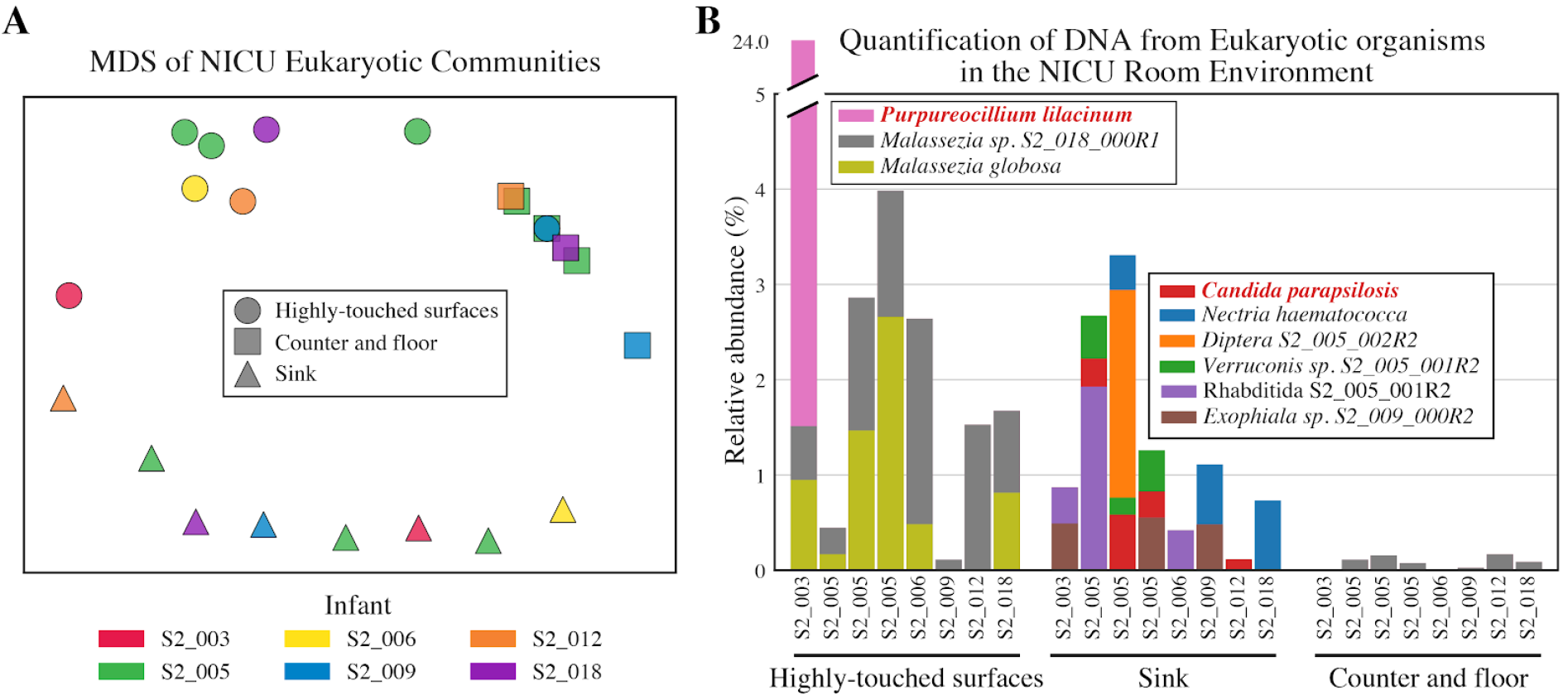
Eukaryotic microbiome of the neonatal intensive care unit (NICU). (A) Multidimensional scaling (MDS) of the Bray-Curtis dissimilarity between all NICU samples. Samples cluster by environment type rather than the room or occupant. (B) Compositional profile of eukaryotic organisms detected in the NICU. Each colored box represents the percentage of reads mapping to an organism’s genome, and the stacked boxes for each sample show the fraction of reads in that dataset accounted for by different eukaryotic genomes in each sample.

The mycobiome of the NICU surfaces is dominated by *Malassezia* (Figure 3B). The eukaryotic organisms found in NICU sinks are distinct from, and more diverse than, those found on surfaces. Sink communities contained *Necteria haematococca, Candida parapsilosis, Exophiala,* and *Verruconis,* all of which were detected in multiple rooms and samples. Additionally, sinks in three separate NICU rooms contain DNA from *Rhabditidia S2_005_000R1* (a novel nematode; see previous section for details). *Diptera S2_005_002R2* (fly) also makes up about 2% of the entire community for single time-point in the sink in infant S2_005’s room (Figure 3B). No macroscopic organisms were noted during the sample collection process. It remains to be seen whether these organisms contribute to dispersal of organisms throughout the NICU or affect the communities themselves.

**Table 2.**
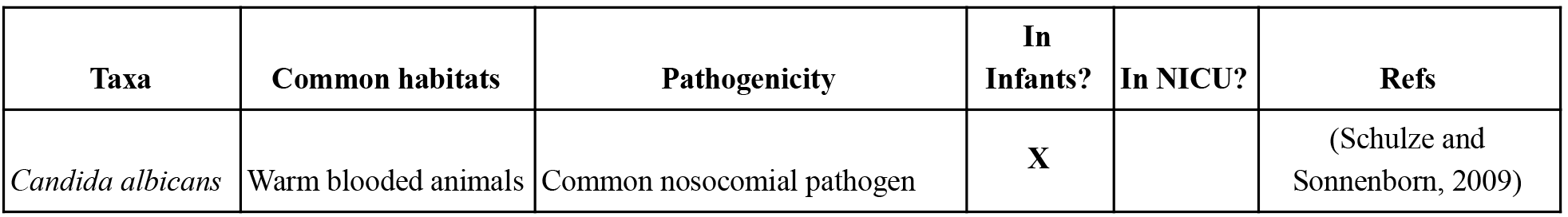
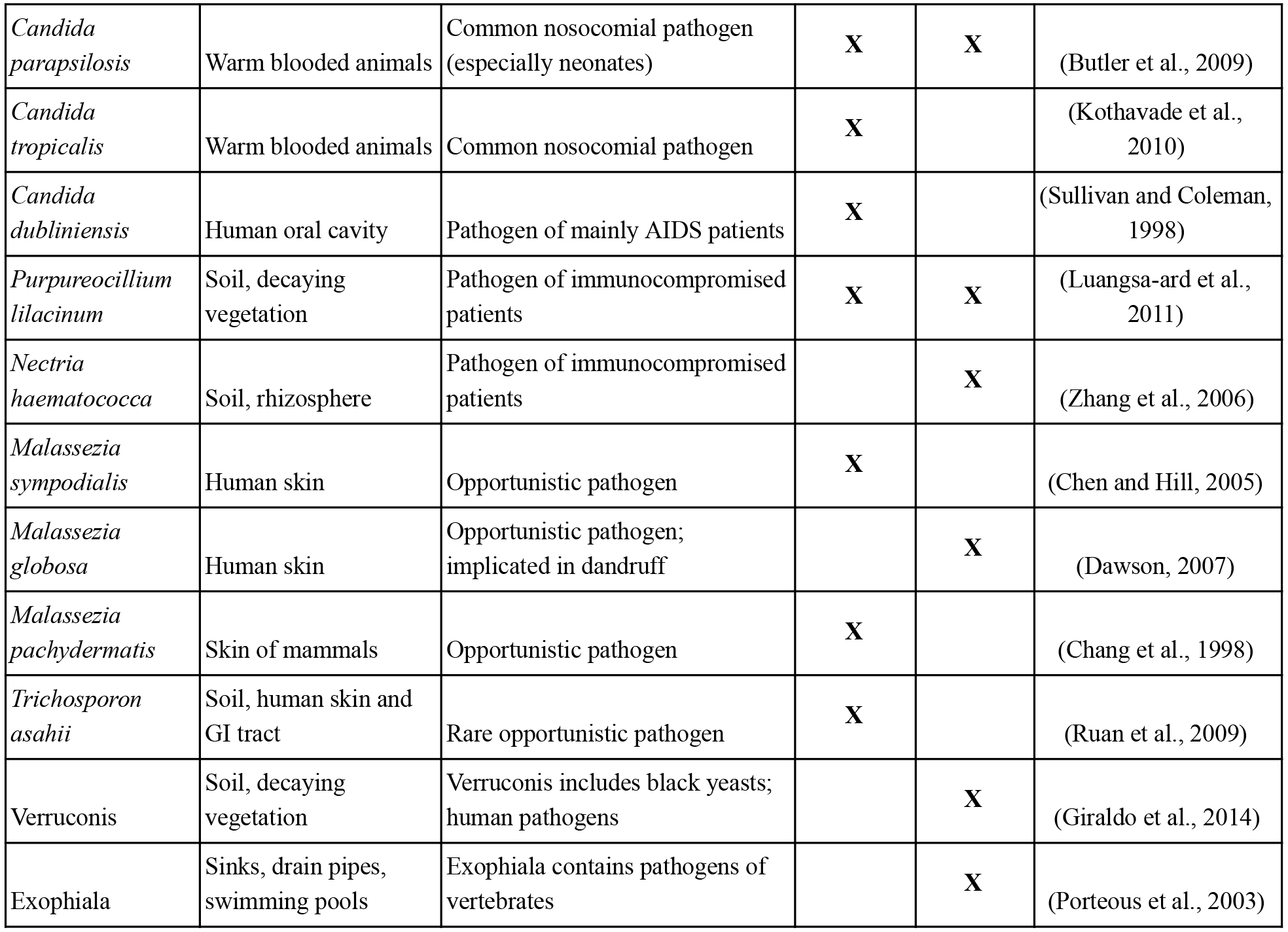
Description of detected fungal taxa.

### Near-identical fungal strain found in NICU and premature infant

Two eukaryotic species were detected in both the NICU and in a premature infant: *Candida parapsilosis* and *Purpurocillium lilacinum.* To contextualize the similarity between them, the genomes assembled from both the infant and room environments were compared to all available reference genomes and each other using dRep (Olm et al., 2017a). *C. parapsilosis* genomes from the NICU sink of infant S2_005 and gut of infant N3_182 were more similar to reference genomes than each other **(Supplemental Figure S1)** whereas *P. lilacinum* genomes from the NICU surfaces of infant S2_003 and gut of infant S2_018 were much more similar to each other than to reference genomes (Figure 4A).

A mapping-based high-resolution genomic comparison was performed to compare the *P. lilacinum* strains in the room and infant (see methods for details). The two genomes have an average nucleotide identity (ANI) of over 99.999%, as well as no detectable differences in genome content (Figure 4B). The NICU sample harboring *P. lilacinum* was collected about 300 days prior to the birth of infant S2_018, and from a different room of the same hospital.

**Figure 4.**
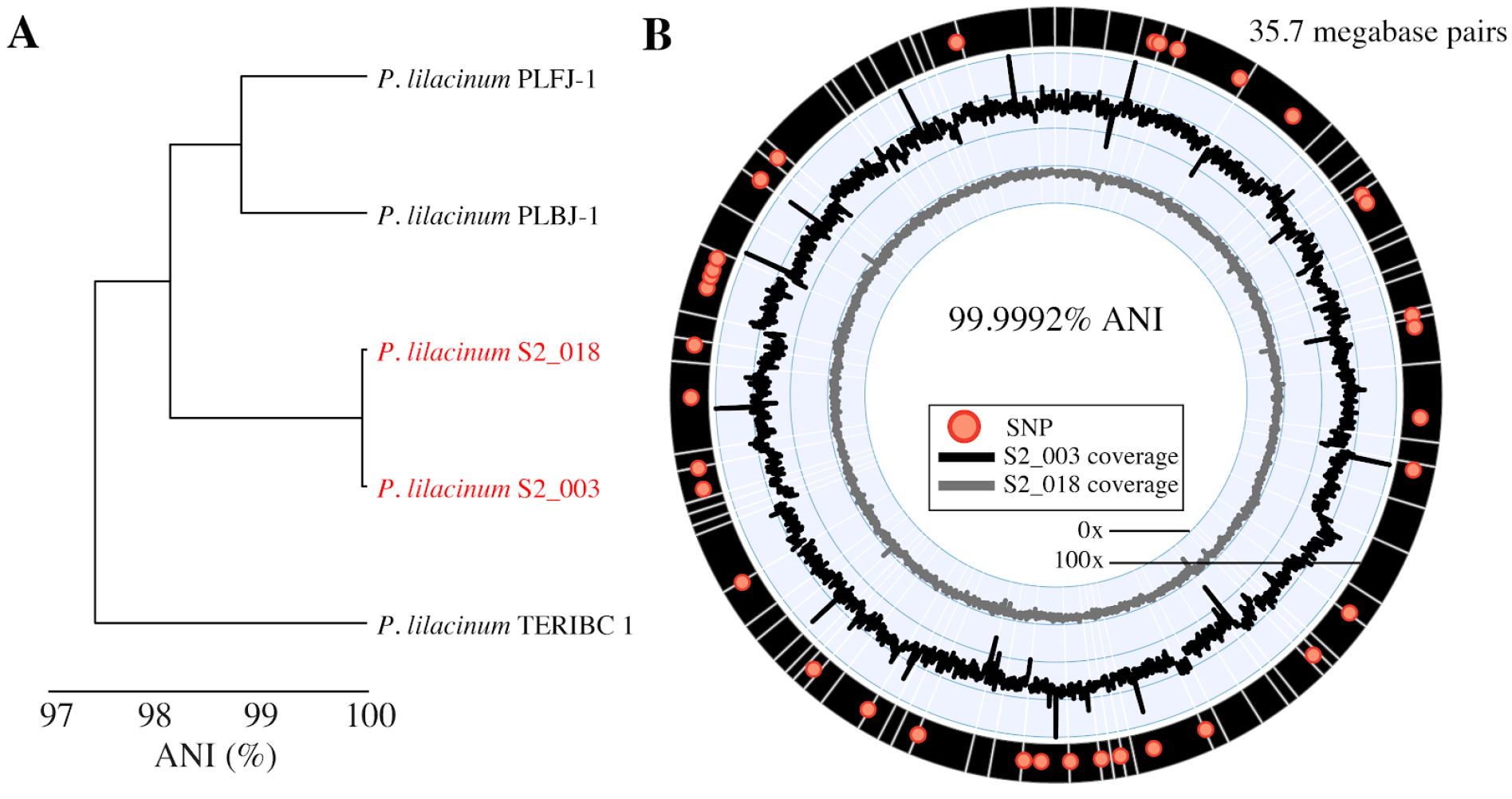
Near identical fungi detected in the NICU and premature infant gut. (A) Dendrogram of *P. lilacinum* genomes based on genome-wide average nucleotide identity (ANI). *P. lilacinum* genomes recovered in this study are red and publicly available reference genomes are black. (B) Detailed genomic comparison of *P. lilacinum* S2_003 and *P. lilacinum* S2_018. For each organism, reads from the sample containing the organism at the highest relative abundance were mapped to the *P. lilacinum* S2_003 genome. Genome-wide coverage is shown in internal line-plots. Scaffolds making up the *P. lilacinum* S2_003 genome are the black rectangles at the edge of the figure, and point-mutations that distinguish the genomes are shown as red dots. Stable genome-wide coverage values suggest that there are no major differences in genome content, and ANI between the genomes was calculated to be 99.9992% based on the small number of detected point-mutations.

### Sequence analysis of new genomes

*De novo* assembly of eukaryotic genomes from metagenomes not only allows for detailed genomic comparison and detection of novel organisms, but also the determination of ploidy, aneuploidy (abnormal number of chromosomes in a cell), heterozygosity, and population heterogeneity of organisms *in vivo.* Changes in ploidy and aneuploidy have been observed in many eukaryotes, especially yeasts (Hirakawa et al., 2015; Peter et al., 2018), and are thought to be a strategy for relatively quick adaptation to shifts in environmental conditions. To determine the ploidy of our genomes, we examined the read count for each allele at a given variant site. For a diploid genome, alleles are expected to have a read count of 50%; for a triploid genome, alleles are expected to have a read count of either 33% or 67%. Based upon this analysis, all of our reconstructed genomes are diploid (Figure 5A, **Supplemental Figure S2**). Similarly, aneuploidy can be detected by searching for regions where allele frequencies and/or read coverage differ from the rest of the genome. We did not see evidence for aneuploidy in any of our genomes **(Supplemental Figures S3, S4)**. However, it is possible regions of aneuploidy may be unintentionally excluded during the binning process due to altered coverage values.

**Figure 5.**
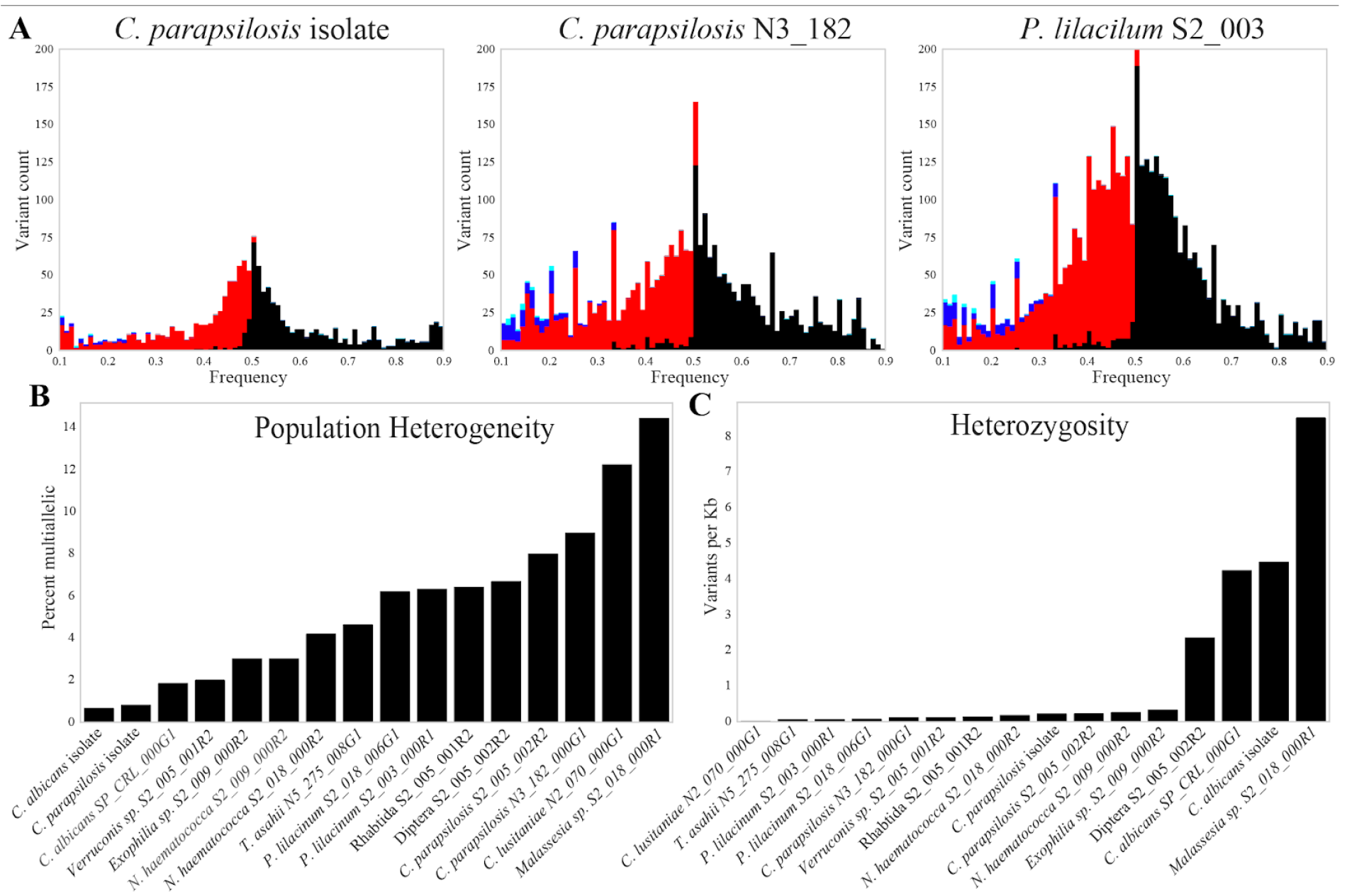
Ploidy and zygosity of recovered eukaryotic genomes. (A) Histogram of the frequencies of the four most abundant variants at each variant site in three genomes. Black, red, dark blue and light blue bars indicate the abundances of the most abundant, second, third and fourth most abundant variant, respectively. (B) For each genome, black bars indicate the percentage of variant sites that are multiallelic (contain more variants at a site than would be expected based upon ploidy alone). (C) For each genome, black bars indicate the number of heterozygous variants per kb across the entire assembled genome.

For diploid genomes reconstructed from metagenomes, the sequences for each chromosome are a composite of sequences from the two alleles. Further, there may be population heterogeneity in the from of additional sequence variants for each allele. Population heterogeneity can be detected based on read counts that exceed the expected ratio (i.e., as multiallelic sites). Measuring population heterogeneity in this way is confounded by sequencing error and stochastic read coverage variation. Genomic datasets for isolates are not expected to have population heterogeneity but will display sequencing error and stochastic read coverage variation. Consequently, we could separate sequencing noise from true population heterogeneity by comparing the patterns we observed in our population genomic data to heterogeneity found in isolate genomic datasets. In our genomes (*C. parapsilosis* N3_182_000G1 and *P. lilacinum* S2_003_000R1), the peak of allele frequencies appears to be much wider than that of the sequenced *Candida parapsilosis* isolate (Figure 5A), suggesting substantial population heterogeneity. Simulated read datasets demonstrate that low coverage can amplify the effect of stochastic read coverage **(Supplemental Figure S5)**, so genomes below 50x coverage were excluded from this analysis. When examining the presence of multiallelic sites, all of our genomes have more multiallelic sites than isolate sequenced genomes (Figure 5B), suggesting that all of our genomes have appreciable population heterogeneity.

Overall heterozygosity for each genome was measured by calculating the number of heterozygous SNPs per kilo-base pair (Figure 5C). A wide range of heterozygosity was observed within genomes. For most organisms, the chromosome copies are similar, and for *C. albicans* and *C. parapsilosis*, comparable to that of reference isolates. *Malassezia* sp. S2_018_000R1 has both a particularly high rate of SNPs per kilo-base pair and high population heterogeneity.

## DISCUSSION

### Eukaryotic genome recovery from metagenomes augments information from isolate studies

In contrast with prior studies that have investigated microbial eukaryote genomes via sequencing of isolates, the current study employed a whole community sequencing approach that enabled evaluation of the diversity of organisms present within populations. For example, prior studies of *C. albicans* have reported that the organism grows clonally *in vivo* (Anderson et al., 2001). However, it is well known that *Candida,* when expressing a certain phenotype, reproduces sexually (Hull et al., 2000). Thus, it is expected that a natural population present within a human or room microbiome would undergo sexual reproduction with distinct coexisting strains, leading to the production of new genotypes. In the current analysis, most of the samples contained one dominant eukaryotic genotype, presumably one well adapted to the habitat, but other allele variants indicate the presence of lower abundance genotypes (Figure 5B). We detected similar levels of population heterogeneity in the NICU and infant-derived samples. In the case of sinks, the reservoir is intensively cleaned on a regular basis so it seems likely that the organisms found there are selected based on their colonization ability. Similarly, the infant gut is near-sterile at the time of birth so organisms found there may be the specific genotypes well suited to be colonists.

Because all populations were dominated by one genotype, it was possible to directly determine that most genomes have relatively low heterozygosity, indicating diversity reduction due to inbreeding and/or strong selection for specific alleles. We speculate that this reflects a long history of colonization of a habitat type that strongly selects for a specific genotype, so genome structure reflects the relatively low probability of recombination with strains with divergent alleles (in other words, the presence of gut-adapted and sink-adapted strains). However, without the availability of similar genomes to compare to from other habitats, we cannot rule out genetic bottlenecks that took place prior to introduction to the hospital.

Four strains were determined to have relatively high heterozygosity, similar to that observed in previously sequenced isolate genomes. *Malassezia* on NICU surfaces has particularly high heterozygosity and population heterogeneity. *Malassezia* are skin associated fungi (Gaitanis et al., 2012), so their high heterozygosity and population heterogeneity may be the consequence of the accumulation of numerous strains throughout the hospital via skin shedding. This could also reflect naturally large population variation present within the skin of a single individual, as has been reported for skin-associated bacteria (Oh et al., 2014; Tsai et al., 2016).

### Eukaryotes in the NICU have the genomic potential to colonize hospitalized infants

The presence of essentially identical *Purpureocillium* strains in an infant gut and a room surface sample collected almost a year apart could reflect a sustained, widespread population of *P. lilacinum* in the NICU. Persistence could be achieved through long-term carriage by a healthcare worker, continual passage between infants and rooms, or a combination of these factors. We also considered the possibility that these sequences represent contaminants, possibly introduced during sequencing. Arguing against this, *P. lilacinum* is known to colonize immunocompromised patients, and the strains in the gut samples display same time series abundance patterns of other gut-associated fungi (Figure 2A). Further, the samples from the same infant were all processed at the same time using the same reagents, yet abundances change substantially across the sample series. Additionally, the two *Purpureocillium* genomes reconstructed from gut samples from different infants are distinct (Figure 4); one would expect the sequences to be more similar if derived from the same reagent contaminants (Olm et al., 2017b).

*C. parapsilosis* was identified in two premature infants and four NICU samples whereas *C. albicans* was identified in six premature infants but not identified in NICU samples. This is interesting because previous studies have linked neonatal *C. albicans* acquisition to vertical maternal transmission, but were unable to identify the source of *C. parapsilosis (Waggoner-Fountain et al., 1996).* Although different strains of *C. parapsilosis* were found in the NICU and in infants, the species-level detection points to the NICU as a possible reservoir for *C. parapsilosis* strains that colonize infants.

All fungal species detected in the infant and NICU environments have been previously implicated as pathogens of immunocompromised individuals (Table 2). Some species, like *Candida albicans,* are almost exclusively found in warm-blooded animals and are thus adapted to growth in humans. Other species, like *P. lilacinum*, are usually associated with soil and decaying plant matter. However, previous studies have shown that strains that colonize humans are similar to environmental strains (Luangsa-ard et al., 2011). Thus, the presence of these strains in the NICU environment could pose risks to these immunocompromised premature infants.

### Premature infants are colonized by eukaryotes early in life

We compared our findings related to premature infant colonization by fungi to three previous studies of fungal colonization of premature infants, one of which used culturing-based detection (Baley et al., 1986), one denaturing gradient gel electrophoresis (DGGE) (Stewart et al., 2013) and the third high-throughput ITS sequencing (LaTuga et al., 2011). The infant colonization rates of these studies were 26%, 50%, and 63%, respectively, thus highly variable and substantially higher than the colonization rate noted here (8%). Compared to shotgun sequencing, DGGE and ITS methods should be more sensitive due to the use of PCR. However, the ability to amplify very rare sequences from organisms present at exceedingly low abundance levels complicates interpretation of the measured colonization frequencies. Counter to this, the shotgun sequencing-based methods provide a more balanced view of community composition (because the methods do not rely on PCR). However, sequencing allocations used in this study (median 3.5 giga-base pairs / sample) typically enable detection only of more abundant populations (e.g., 0.35 % of the community DNA; **Supplemental Table S1)**. It is not known whether the seemingly highly restricted period of colonization by eukaryotes is due to low sensitivity of the shotgun sequencing method.

However, it is also possible that this is a general phenomenon, as most other studies have not analyzed finely resolved time-series data as used in this study.

All infants profiled in this study received 2-7 days of prophylactic antibiotics upon birth, meaning antibiotic use is highly correlated with earlier days of life **(Supplemental Table S3)**. While both antibiotic administration and DOL were significantly correlated with eukaryote abundance, infants who received antibiotics later in life were not colonized by eukaryotes, suggesting that day of life is the more important factor. Proteobacteria were also significantly anti-correlated with eukaryote abundance. This is interesting because Proteobacteria are usually enriched in early life samples and in infants who have received antibiotics (Hu et al., 2016; Rodríguez et al., 2015). Many important pathogens of infants are Proteobacteria, including *Escherichia coli, Klebsiella pneumoniae*, and *Pseudomonas aeruginosa* (McGuire et al., 2004). Future studies are needed to determine if there are important associations of other bacteria with fungi, and if microbial eukaryotes inhibit growth of these pathogens in preterm infants or vice versa.

### Differences in colonization patterns of NICU sinks and surfaces

Yeasts of the genus *Malassezia,* which dominated NICU surfaces, are the most common eukaryotic member of the healthy skin microbiome (only one NICU surface sample contained a eukaryote not from this genus, Figure 3) (Gaitanis et al., 2012; Parfrey et al., 2011). Similarly, previous studies showed that hospital surfaces (and other built environments) are dominated by typically skin-associated bacteria (Brooks et al., 2017, 2018; Chase et al., 2016; Hewitt et al., 2013; Shin et al., 2015). The broad consistency in the eukaryotic communities in surface samples taken in different NICU rooms a year apart could be explained by continuous passive accumulation of healthy skin fungi shed by healthcare workers, punctuated by cleaning events.

The same organisms were never detected in sinks and surfaces. In contrast to surfaces, the sinks hosted a diverse and variable eukaryotic community and sink communities in different rooms were distinct. Although all fungi were detected in at least two sink-associated samples, no eukaryote was present in over half of the samples (Figure 3). The only organism present in a single sample was *Diptera* S2_005_002R2 (fly). As no macroscopic organisms were detected during the collection process, the DNA may have come from sink-associated eggs. Recent studies have suggested that insects play significant roles in the dispersal of fungi, and this may occasionally occur in the NICU (Madden et al., 2018). The other metazoan detected, the worm *Rhabditida* S2_005_001R2, was found in sinks from multiple rooms and collected months apart. These organisms may also be a source of fungi, and like the fly, could impact the overall NICU microbiome.

### Conclusions

We applied genome-resolved metagenomics to study eukaryotes in the gut microbiomes of infants and their NICU rooms. We detected eukaryotes associated with pathogenesis of immunocompromised humans, commensals of human skin, and fungi typical of environments such as soil and drain pipes. Genomic analysis of diploid organisms found low rates of heterozygosity that may be explained by persistence of hospital-associated lineages in environments that impose strong selective pressure. The application of this approach in other contexts should greatly expand what is known about eukaryote diversity, population variation, and dissemination pathways.

## ACKNOWLEDGEMENTS

This research was supported by the National Institutes of Health (NIH) under award RAI092531A, the Alfred P. Sloan Foundation under grant APSF-2012-10-05, and National Science Foundation Graduate Research Fellowships to M.O. and P.W. under Grant No. DGE 1106400. The study was approved by the University of Pittsburgh Institutional Review Board (Protocol PRO10090089). This work used the Vincent J. Coates Genomics Sequencing Laboratory at UC Berkeley, supported by NIH S10 0D018174 Instrumentation Grant. We thank Christopher T. Brown for helpful discussions.

## Data availability

Eukaryotic genomes are available at NCBI BioProject PRJNA471744, as well as at https://github.com/MrOlm/InfantEukaryotes. Newly sequenced infant metagenomes are available at NCBI BioProject PRJNA417343. NICU room metaganomes are available from the Short Read Archive (accessions SRR5420274 to SRR5420297) (Brooks et al., 2017), whereas infant metagenomes are available under Bioproject PRJNA294605, SRA studies SRP052967, SRP114966, and SRP012558, and SRA accessions SRR5405607 to SRR5406014 (Brooks et al., 2017; Brown et al., 2018; Rahman et al., 2018; Raveh-Sadka et al., 2015, 2016; Sharon et al., 2013)). Python Jupyter notebooks detailing figure generation and statistical analyses are available on GitHub https://github.com/MrOlm/InfantEukaryotes.

## Author Contributions

M.O., B.B., M.J., and J.F.B. conceived of study design. M.O. and P.W. performed computational analysis. R.B. recruited study subjects and collected DNA samples, and B.F. performed DNA extractions. M.O., P.W., and J.F.B. wrote the manuscript, and all authors contributed to manuscript revisions.

## METHODS

### Subject recruitment, sample collection, and metagenomic sequencing

This study was reviewed and approved by the University of Pittsburgh Institutional Review Board (IRB PRO12100487 and PRO10090089). This study made use of many different previously analyzed infant datasets. These datasets have previously published descriptions of the study design, patient selection, and sample collection, and are referred to as NIH1 (Brown et al., 2018; Raveh-Sadka et al., 2016), NIH2 (Brooks et al., 2017), NIH3 (Raveh-Sadka et al., 2015), NIH4 (Rahman et al., 2018), Sloan2 (Brooks et al., 2017), and SP_CRL (Sharon et al., 2013).

This study also involved the collection and processing of an additional 269 samples from 53 infants. Newly collected infant fecal samples followed the same sample collection and DNA extraction protocol as described previously (Olm et al., 2017c). Metagenomic sequencing of newly collected infant fecal samples was performed in collaboration with the Functional Genomics and Vincent J. Coates Genomics Sequencing Laboratories at the University of California, Berkeley. Library preparation on all samples was performed using the following basic protocol: 1) gDNA shearing to target a 500bp average fragment size was performed with the Diagenode Bioruptor Pico, 2) end repair, A-tailing, and adapter ligation with an Illumina universal stub with Kapa Biosystems Hyper Plus Illumina library preparation reagents, 3) a double AMpure XP bead cleanup, followed by indexing PCR with dual-matched 8bp Illumina compatible primers. Final sequence ready libraries were visualized and quantified on the Advanced Analytical Fragment Analyzer, pooled into 11 subpools based on mass and checked for pooling accuracy by sequencing on Illumina MiSeq Nano sequencing runs. Libraries were then further purified using 1.5% Pippin Prep gel size selection assays collecting library pools from 500-700bp. Pippins pools were visualized on fragment analyzer and quanted with Kapa illumina library quant qPCR reagents and loaded at 3nM. The eleven pools were then sequenced on individual Illumina HiSeq4000 150 paired-end sequencing lanes with 2% PhiX v3 spike-in controls. Post-sequencing bclfiles were converted to demultiplexed fastq files per the original sample count with Illumina’s bcl2fastq v2.19 software. New metagenomic data was processed in the same manner as in the prior studies, and as described in Rahman 2018.

### Eukaryotic genome binning and gene prediction

Binning assembled sequence scaffolds into eukaryotic genomes was performed using a EukRep based pipeline, described in detail in West et al. 2018. Given the sequencing depth performed in this study, we calculated that genomes present at over 1% of the population should achieve the 8x coverage typically needed for genome assembly (**Supplemental Figure S6**). In cases where time-series data were available, samples were pre-binned using time-series information and eukaryotic bins were then subsequently identified with EukRep. In addition to the gene prediction methodology outlined previously (West et al., 2018), a second homology-based gene prediction step was performed. Ribosomal S3 (rpS3) proteins were identified in genomes using a custom ribosomal protein S3 (rpS3) profile HMM, and identified sequences were searched against the NCBI database (NCBI Resource Coordinators, 2017) and UniProt (UniProt Consortium, 2015) using blast (Altschul et al., 1990). For each *de novo* assembled genome, gene sets for the top 1-3 most similar organisms were used as homology evidence for a second pass gene prediction step with AUGUSTUS (Stanke et al., 2006), as implemented in MAKER (Cantarel et al., 2008). For *Rhabditida* S2_005_001R2 first pass gene predictions were used, as homology evidence decreased overall estimated genome completeness.

To verify bins, the taxonomy of each scaffold was determined by searching gene sequences against the UniProt database (Raveh-Sadka et al., 2015). All bins were found to have a consistent phylogenetic signal, except the bin created from sample S2_009_000R2. Scaffolds had similar GC content and sequencing coverage, but were either dominated by genes with homology to the Class Sordariomycetes or Eurotiomycetes. Scaffolds from the original “megabin” were split into two seperate bins based on this phylogenetic signal, resulting in the genomes *Nectria haematococca* S2_009_000R2 and *Exophiala sp.* S2_009_000R2. Gene prediction was run again for both of these genomes, as described above.

### Phylogenetic Analyses

In order to construct a phylogenetic tree, *rpS3* proteins from each *de novo* genome were detected as described above and searched against the NCBI database using blast. Protein sets of the 3-5 most similar organisms on NCBI were downloaded for inclusion. Other phylogenetically important genomes, such as *A. thaliana,* were included as well. For each protein set, 16 ribosomal proteins (bacterial ribosomal protein names L2, L3, L4, L5, L6, L14, L15, L16, L18, L22, L24, S3, S8, S10, S17, and S19) were identified using custom built hidden markov models (HMMs) with HMMER (Finn et al., 2011), using the noise cutoff (NC). The 16 ribosomal protein datasets were then aligned with MUSCLE (Edgar, 2004) and trimmed by removing columns containing 90% or greater gaps. The alignments were then concatenated. A maximum likelihood tree was constructed using RAxML v.8.2.10 (Stamatakis, 2006) on the CIPRES web server (Miller et al., 2010) with the LG plus gamma model of evolution (PROTGAMMALG) and with the number of bootstraps automatically determined with the MRE-based bootstrapping criterion. The constructed tree was visualized with Interactive Tree of Life (ITOL) (Letunic and Bork, 2007).

Average nucleotide identity (ANI) between binned genomes and reference genomes was determined with dRep (Olm et al., 2017a). Resulting whole genome ANI values were used in combination with a 16 ribosomal protein phylogenetic tree to determine the taxonomy of *de novo* genomes. For genomes without a genus-level taxonomy, mitochondrial *COI* genes were identified by searching *D. melanogaster* and *C. elegans COI* genes against our PRODIGAL (Hyatt et al., 2010) predicted genes sets with UBLAST (Edgar, 2010). Significant hits from our protein sets were then searched against the Barcode Of Life Database (BOLD) (Ratnasingham and Hebert, 2007) and NCBI in order to identify sequences with high identity to our novel genomes.

### Mapping-based genome detection

To detect eukaryotes in an assembly-free manner, reads were mapped to a curated genome collection. This genome collection consists of all fungal genomes in RefSeq (accessed 9/14/17) (Pruitt and Maglott, 2001), as well as genomes assembled in this study with no close representatives in RefSeq (average nucleotide identity of 90% or higher according to Mash (Ondov et al., 2016)). The six genomes with no close representatives in RefSeq were *Malassezia sp.* S2_018_000R1, Diptera S2_005_002R2, *Exophiala sp.* S2_009_000R2, *Verruconis sp.* S2_005_001R2, and Rhabditida S2_005_001R2. *Candida parapsilosis* CDC317 was also included, as there were no genomes of *C. parapsilosis* in RefSeq.

Reads from all samples were mapped to this reference genome list using Bowtie 2 (Langmead and Salzberg, 2012). An organism was considered present in a sample if it had a breadth of coverage of at least 10%, as determined by pipe_coverage (https://github.com/MrOlm/pipeCoverage). The lowest coverage genome with this breadth threshold was 0.35x coverage. In order to determine the limit of detection, we determined relative abundance needed to achieve 0.35x coverage using the median infant sequencing depth (3.5 giga-base pairs) and the median eukaryotic genome length in our database of organisms that were detected at least once. This lead to an estimated limit of detection of 0.35% relative abundance.

### Statistical analyses and generation of MDS plot

To compare the euakryotic communities present in NICU room samples, multidimensional scaling (MDS) based on Bray-Curtis distance was performed. Bray-Curtis distance was calculated based on the relative abundance of each eukaryote in present a sample using the python library SciPy (command scipy.spatial.distance.braycurtis) (Jones et al., 2001). Eukaryotes with at least 10% breadth of coverage were considered present in a sample. MDS was performed on the resulting all-vs-all distance matrix using the python library sklearn (command sklearn.manifold.MDS) (Pedregosa et al., 2011). MDS was plotted using a custom function in Matplotlib (Hunter, 2007). Open source code detailing this analysis available at https://github.com/MrOlm/InfantEukarvotes.

We tested for significant associations between samples containing eukaryotes and various forms of metadata using the python SciPy package (Jones et al., 2001). Included were six pieces of continuous metadata (DOL, infant birth weight, ect.), twenty-three pieces of categorical metadata (specific antibiotics given and specific NICU room locations), and the phyla-level abundance of all bacterial genomes (seven total phyla) **(Supplemental Table S3)**. Bacterial phyla-level abundance was determined by summing the relative abundance of all bacterial genomes present in a sample. Bacterial genomes for previously sequenced samples are available in a previous publication (Rahman et al., 2018), and bacterial genomes for newly sequenced genomes were binned using the same methods. Metadata was filtered such that between 20-80% of values were non-zero in both samples containing eukaryotes and samples not containing eukaryotes. This resulted in a total of 13 pieces of meta-data for statistical testing **(Supplemental Table S3)**.

In order to eliminate statistical bias introduced through sampling the same infant multiple times, one sample from each infant was chosen for statistical tests. If the infant was not colonized by a eukaryote, the sample was chosen at random. If the infant was colonized by a eukaryote, the sample with the highest eukaryotic abundance was chosen. Samples were considered to have a eukaryote present if the sum of the relative abundance of eukaryotes with at least 10% breadth was at least 0.1% relative abundance. Fisher’s exact test was used for categorical metadata, and Wilcoxon rank-sum test was used for continuous data. Benjamini-Hochberg p-value correction (Benjamini and Yekutieli, 2001) was performed to account of multiple hypothesis testing. The results of all statistical tests are provided in **Supplemental Table S4**. Open source code detailing this statistical analysis available at https://github.com/MrOlm/InfantEukaryotes.

### Detailed genomic comparisons

dRep v1.4.3 (Olm et al., 2017a) was used to generate the dendrograms presented in Figure 4A and **Supplemental Figure S1**, using the command “dRep cluster --SkipMash”. MUMmer (Delcher et al., 2003) was used to align scaffolds from genome *Purpureocillium lilacinum* S2_018_006G1 (query) to genome *Purpureocillium lilacinum* S2_003_000R1 (reference), as implemented in dRep using the method “ANImf”. The resulting alignment was then filtered in a number of ways. Only the best alignment from each query scaffold to the reference was retained, alignments with less than 1 kilo-base pair gaps were merged, in cases where multiple query sequenced aligned to the same reference sequence only the longest alignment was retained, and all alignments less than 100 base pairs were removed. These filtering steps reduced the total length of aligned sequence by 3%. Aligned scaffolds were depicted on the outer ring of Figure 4B using circos (Krzywinski et al., 2009).

Point mutations between both recovered *Purpureocillium lilacinum* genomes were detected by mapping reads from all samples from infant S2_003 and the NICU metagenome of infant S2_018 to genome *Purpureocillium lilacinum* S2_003_000R1 using Bowtie 2 (Langmead and Salzberg, 2012). The coverage of these alignments was calculated for 10 kilo-base pair windows using pipe_coverage (https://github.com/MrOlm/pipeCoverage). PileupProfile was used to identify SNPs from the resulting alignment (https://github.com/banfieldlab/mattolm-public-scripts/blob/master/PileupProfile.py). Under the default settings used, SNPs are called in locations with at least 80% of reads differing from the genome base call, and only at locations that are at least 50 base pairs from the edge of a scaffold. SNPs were called using reads from both infants, and SNPs at locations from both read sets were discarded. All remaining detected SNPs were depicted on the outer ring of Figure 4B using circos (Krzywinski et al., 2009).

### Ploidy, heterozygosity, and population heterogeneity

In order to identify variants, reads from the sample a particular genome was binned from were mapped back to the *de novo* assembled genome using Bowtie2 (Langmead and Salzberg, 2012) with default parameters. The PicardTools (http://broadinstitute.github.io/picard/) functions “SortSam” and “MarkDuplicates” were used to sort the resulting sam file and remove duplicate reads. Freebayes (Garrison and Marth, 2012) was used to perform variant calling with the options ‘--pooled-continuous −F 0.01 −C 1’. Variants were filtered downstream to include only those with support of at least 10% of total mapped reads in order to avoid false positives. SNP read counts were calculated using the ‘AO’ and ‘RO’ fields in the freebayes vcf output file. Multiallelic sites were defined as sites with two or more non-reference alleles. Variants were called using the same methodology for both simulated read datasets and isolate genomes. Variants were used to determine ploidy, heterozygosity, and population heterogeneity as described in the results section. Source code with full implementation details available at https://github.com/MrOlm/InfantEukarvotes.

In order to determine the effect of stochastic read coverage on variant frequencies, simulated haploid, diploid, and triploid genomes were generated using the pIRS (https://github.com/galaxy001/pirs) diploid command with the *C. albicans* P57072 reference genome. The command was used once to generate a diploid genome, and twice to generate a triploid genome. Simulated reads were then generated for each genome using the pIRS simulate command at 10x, 50x, and 100x coverage. Assemblies and raw reads were downloaded for both *C. albicans* A48 and *C. parapsilosis* CDC317 from NCBI to be used as example isolate genomes for comparison. Based on this analysis, genomes below 50x coverage were excluded from this analysis.

Genome aneuploidy was analyzed in two ways. First, reads from each sample were mapped back to genomes assembled from that sample. The coverage of each scaffold was determined in 10 kilo-base pair windows, and the coverage of all windows for each scaffold over 10 kilo-base pairs was plotted. Plots were then analyzed for scaffolds with differing coverage, indicative of the presence of multiple copies. **(Supplemental Figure S3)**. Second, reads from samples with genomes assembled from them were mapped to the closest available reference genome. The same procedure was then performed with these reference genomes in all cases where at least 80% of the genome was covered by reads. This allowed determination of aneuploidy on the whole-chromosome level **(Supplemental Figure S4)**. Both methods agreed in all cases, and no aneuploidy was detected.

## SUPPLEMENTAL INFORMATION

Available at https://github.com/MrOlm/InfantEukaryotes

**Supplemental Figure S1.** Dendrogram of reference and *de novo* assembled *C. parapsilosis* genomes based on genome-wide average nucleotide identity (ANI).

**Supplemental Figure S2.** Raw variant frequency graphs used to determine ploidy of all *de novo* assembled genomes.

**Supplemental Figure S3.** Determination of aneuploidy for all *de novo* assembled genomes.

**Supplemental Figure S4.** Alternative determination of aneuploidy for genomes with high quality reference genomes.

**Supplemental Figure S5.** Effect of coverage on variant frequency determination as assessed through simulation of metagenomic reads.

**Supplemental Figure S6.** Evaluation of relative abundance, genome length, and sequencing depth needed to achieve 8x sequencing coverage.

**Supplemental Table S1.** Sequencing metadata for all infant and room metagenomic samples

**Supplemental Table S2.** Mapping-based abundance of eukaryote genomes in all samples

**Supplemental Table S3.** Metadata for statistical associations

**Supplemental Table S4.** Statistical associations of samples containing eukaryotes with metadata.

